# A single UV-C pulse modulates Gibberellin homeostasis and Plant Development in Arabidopsis

**DOI:** 10.64898/2026.04.28.721437

**Authors:** Maria J. Pimenta Lange, Adolfo Calvo-Parra Martínez, Johanna Schwarze, Theo Lange

## Abstract

Under natural growth conditions, plants are not usually exposed to the high-energy ultraviolet C range (UV-C, 100–280 nm) of the solar spectrum, as this is absorbed by the ozone layer. However, low doses of UV-C radiation can trigger stress responses in plants. Nevertheless, it is not yet fully understood how UV-C light affects plant development at the hormonal level. Here we show that a single one-min UV-C light pulse (20 W/m^2^) alters gibberellin (GA) homeostasis in Arabidopsis in two phases: initially, the level of GA_12_ ‒ a key precursor of the final part of gibberellin biosynthesis ‒ is reduced. Consistent with this, the transcript levels of the *CPS, KS* and *KAO2* genes, which encode enzymes involved in the initial parts of gibberellin biosynthesis, decrease. The level of the plant hormone GA_4_ also decreases initially, probably due to the reduced GA_12_ precursor levels. However, in a second phase, the endogenous GA_4_ levels rise in UV-C treated plants relative to control plants. This increase leads to an early onset of flowering, as well as increased growth and fertility, in UV-C-treated Arabidopsis plants. The GA signalling mutant *gdella* does not exibit wild-type phenotypic responses to UV-C treatment, indicating that GA signalling is essential for the UV-C response. To further narrow down the responsible steps in the GA-signalling pathway, we tested the *kao1* and *kao2* mutants, which are both impaired in early gibberellin biosynthesis. Neither mutant displays phenotypic responses to the UV-C treatment, indicating that both genes are required for mediating the UV-C response. In contrast, the quintuple 2-oxidase mutant C_19-_-*2oxqM* exhibits responses to UV-C treatment similar to the wild-type, suggesting that the five catabolic 2-oxidases that act on C_19_-GAs play a negligible role in regulation GA-hormone levels for growth and development in this case.

**Highlight:** UV-C pulse triggers biphasic gibberellin dynamics, delaying early development but ultimately enhancing growth and fertility in *Arabidopsis thaliana*.

## Introduction

In nature, plants need sunlight that provides energy for growth and guides their development. The ultraviolet (UV) part of the solar spectrum is divided into UV-C (100-280 nm), UV-B (280-315 nm) and UV-A (315-400 nm). UV light accounts for about 7% of the total solar radiation reaching the earth’s surface, and the high-energy radiation UV-C and UV-B are generally considered to be completely and mostly absorbed by the ozone layer, respectively (Tossi *et al*., 2019). However, the UV-C spectral region at the Earth’s surface has recently been reported, challenging current atmospheric ozone models (Herndon *et al*., 2018). Depending on the dose, UV radiation can have both beneficial and harmful effects to plants. If environmental light delivers too much energy to plants, exceeding their ability to use or dissipate it, it can induce stress in plants. It is generally accepted that plants are not affected by UV-C light under natural growing conditions. However, low doses of UV-C irradiation have been used to study stress responses in seed plants (Yao *et al*., 2011; Takeno, 2016) and to enhance plant defence against diseases (Urban *et al*., 2018; Vàsquez *et al*., 2020; Otake *et al*., 2021).

In this study, we investigated how varying durations of UV-C treatment affect growth and development of Arabidopsis plants and how this is regulated by the plant hormone gibberellin. We found that a one-minute UV-C pulse of 1200 J/m^2^ significantly accelerated flowering, increasing plant height and fertility compared to untreated plants. Gibberellins (GAs) are plant hormones known to promote plant growth and development, including flowering (Pimenta Lange and Lange, 2006; Lange and Pimenta Lange, 2020; Hedden, 2020). Gibberellin biosynthesis is complex and, in Arabidopsis, mainly proceeds *via* the non-hydroxylation pathway (Fig. 1). The initial part of the anabolic pathway begins with geranylgeranyl pyrophosphate (GGPP), which is formed in the isoprenoid pathway. Several subsequent steps, catalysed by four enzymes (*ent*-copalyl pyrophosphate synthase (CPP), *ent*-kaurene synthase (KS), *ent*-kaurene oxidase (KO) and *ent*-kaurene acid oxidase (KAO), then form the central precursor GA_12_ of the final part of gibberellin biosynthesis. GA_12_ is a gibberellin with 20 carbon atoms (C_20_-GA) that is ultimately converted into the plant hormone GA_4_ (a C_19_-GA) with the help of GA 20-oxidases and GA 3-oxidases. GA_4_ can be degraded into GA_34_ with the help of GA 2-oxidases (C_19_-2ox). However, degradation can also occur at earlier stages. For instance, GA_110_ is formed from GA_12_ with the help of C_20_-2ox, and GA_25_ is produced from GA_24_ *via* a specific 20ox (see Fig. 1).

**Figure 1.**
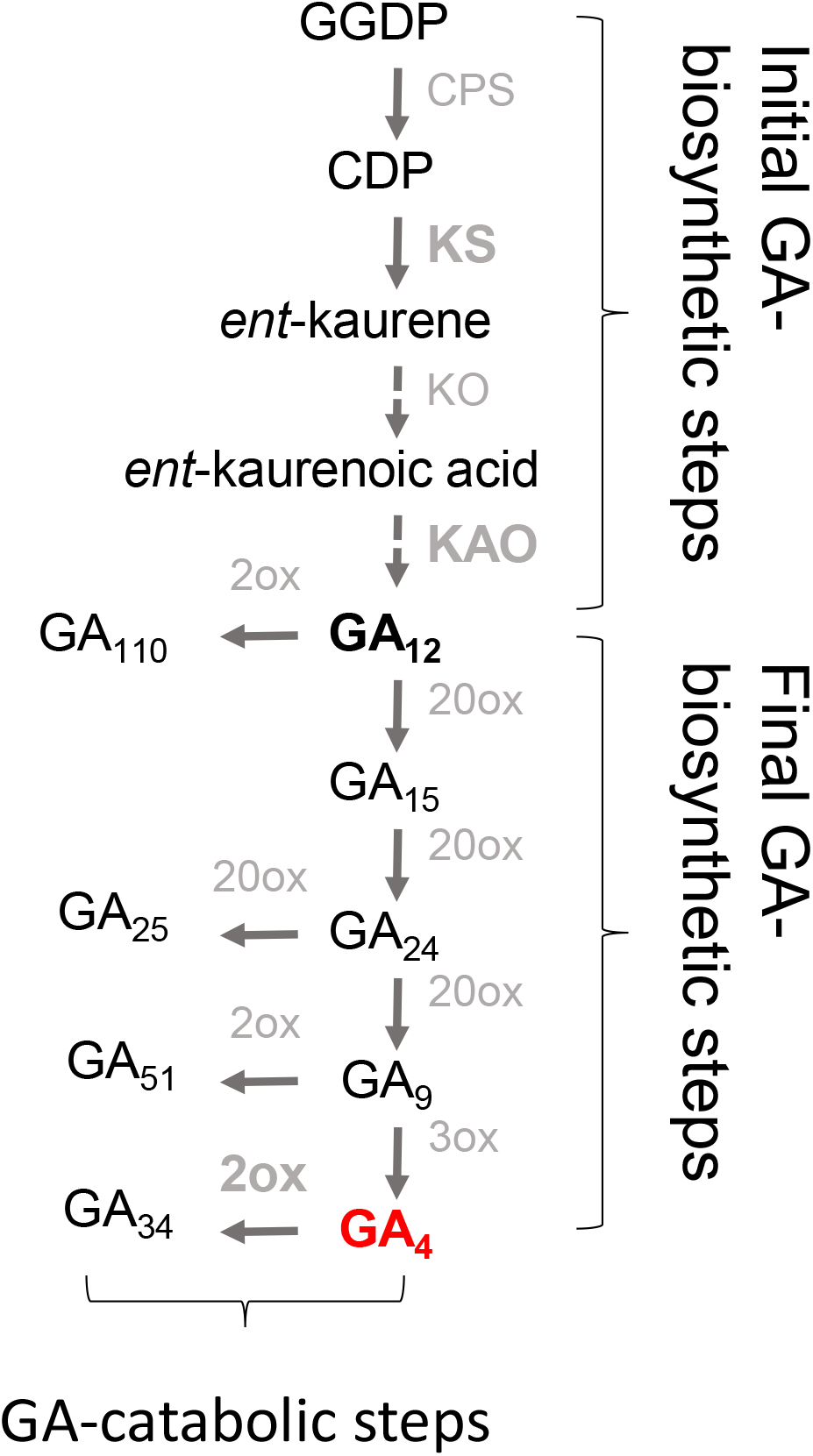
Simplified ‘non-hydroxylation’ gibberellin pathway in Arabidopsis. Enzymes are labelled in grey; CPS, *ent*-copalyl pyrophosphate synthase; KS, *ent*-kaurene synthase; KO, *ent*-kaurene oxidase; KAO, *ent*-kaurenoic acid oxidase; 20ox, gibberellin 20-oxidase; 3ox, gibberellin 3-oxidase; 2ox, gibberellin 2-oxidase. Central precursor GA_12_ and bioactive GA_4_ are highlighted in bold characters.

Here, we demonstrate that UV-C treatment alters GA-biosynthesis in Arabidopsis in two phases. Shortly after a one-minute UV-C pulse, the levels of the GA precursor GA_12_ and of the plant hormone GA_4_ decrease. However, in a second phase, GA_4_ subsequently increases, promoting growth and development. Furthermore, GA signal transduction is crucial for the phenotypic responses to UV-C; the *gdella* mutant, which has impaired GA signal transduction due to the absence of all five DELLA proteins, does not show phenotypic responses to UV-C treatment.

The two GA biosynthesis mutants, *kao1* and *kao2*, which are both involved in the initial stages of GA biosynthesis, also fail to display phenotypic responses to UV-C treatment. This suggests that the initial part of the GA-pathway is particularly important for mediating the UV-C response (Fig. 1). These results enhance our understanding of the hormonal mechanisms through which plants respond to UV-C treatment and may be exploited to optimise plant growth and development, thereby contributing to more sustainable and environmentally friendly plant production systems by reducing the need for chemical plant growth regulators.

## Materials and methods

### Plant materials and growth conditions

Arabidopsis Columbia (Col-0) and Landsberg erecta (Ler) were used as wild type ecotypes. The global della (*gdella, rga-t2 gai-t6 rgl1-1 rgl2-1 rgl3-4*) mutant was provided by Dr. Patrick Achard (IBMP, Strasbourg, France) and is in the Ler background (Koini *et al*., 2009). The double mutant *sid2/NahG* (*sid2/N*) was provided by Dr. Pepe Cana Quijada (UMA-CSIC, Málaga, Spain) and is in the Col-0 background (Rosas-Díaz *et al*., 2017). The C_19_-2ox quintuple mutant (*2ox1, 2ox2, 2ox3, 2ox4, 2ox6*) is in the Col-0 background and was a gift from Dr. Andy Philipps (Rothamsted Research, UK). Seeds were sown on soil:vermiculite (2:1 V/V), 80 g per pot, 92% relative field capacity. Plants were grown in chambers with 70% relative humidity and a 16-h light/8-h dark cycle (long-days, LD) or 8-h light/16-h dark cycle (short-days, SD) at 22°C (lights on) and 20°C (lights off). After sowing and during the first week, the light intensity was 120 µmol m^−2^s^−1^ (Osram cool-white fluorescence lamps). After that the plants were moved to a bigger growth chamber with light intensity 200 µmol µmol m^−2^s^−1^ (Osram Powerstar HQI-T 400W/D daylight lamps). Plants were watered three-times per week by adding the necessary amount of water to keep a similar weight in all pots.

### UV-C light treatments and phenotype analysis

Two-week old plants grown in LD or three-week old plants grown in SD were irradiated for 1 min, or otherwise indicated, with UV-C light, 254nm, 1200 J/m^2^ (20 W/m^2^) in a cross-linker. The UV-C irradiated and non-irradiated plants were place in trays using a chessboard muster to avoid growth effects due to the placement. After UV-C irradiation, the SD grown plants were kept one more week under SD and then moved to LD to induce flowering. Respective control (untreated) plants were grown in the same way.

For the phenotype analysis, flowering time was measured when the inflorescence reached 1cm length. The Rosette leaves diameter and plant height (the height of the main inflorescence) of control and UV-C treated plants were measured 10 days after treatment or as indicated in the text. The number of seeds per plant were calculated by total number of siliques x seed number per silique (minimum of 15 siliques).

### Quantitative analysis of endogenous GAs

Quantitative analysis of endogenous GAs has been described before (Lange *et al*., 2020). Fresh plant material was harvested, frozen in liquid N_2_ and ground. GA analysis and gene expression analysis were done from the same ground material.

### Gene expression analysis

Total RNA extraction has been described before (Pimenta Lange and Lange, 2015). DNase I-treatment of total RNA was done using the RapidOut DNA removal kit following the manufactor’s procedure (Thermofisher). First-strand cDNA was synthesized using 300 ng of DNase I-treated total RNA and the SuperScript IV VILO Master Mix in a 5-µL reaction volume according to the manufacturer’s protocol (Thermofisher).

The obtained cDNA was diluted 10 times with nuclease-free water and used in 10 µL RT-qPCR reactions as previously described (Pimenta Lange and Lange, 2015) but with the addition of 5 µL of qPCRBIO SyGreen No-Rox mix (PCR Biosystems). The At2g28390 (*SAND*) gene was used as the internal reference gene. The Real-time PCR reactions were performed using a FastGene Real-Time PCR cycler (Nippon Genetics). Two technical qPCR replicates were done and the relative expression of each gene was calculated using the PCR Miner program (http://www.miner.ewindup.info/) as the average of two replicates. Technical replicates with a difference from the mean of cycle threshold (Ct) higher than 0.5 were excluded. All experiments were done with at least three biological replicates.

The samples were tested for the absence of genomic DNA by qPCR using RNA as template. The primers used for qPCR analysis were described previously (Pimenta Lange and Lange, 2015; Lange *et al*., 2020).

### RNA sequencing and data analysis

Prior to sequencing, the quantity and quality of DNase I-treated total RNA was analyzed by Qubit fluorometer (Thermo Fisher Scientific) and a fragment analyzer (Agilent). The library preparation and strand-specific RNA-sequencing with PolyA selection was done by GENEWIZ Germany (Leipzig). The sequencing was done in an Illumina NovaSeq™ 6000 platform (Illumina) with 150 bp paired-end reads and a total of 48-72 million reads per sample. Sequence reads were trimmed to remove possible adapter sequences and nucleotides with poor quality using Trimmomatic v.0.36. The trimmed reads were mapped to the Arabidopsis thaliana TAIR10 reference genome available on ENSEMBL using the STAR aligner v.2.5.2b. Unique gene hit counts were calculated by using feature Counts from the Subread package v.1.5.2. The hit counts were annotated using the gene-id. Data quality assessments were performed to detect any samples that are not representative of their group, and thus, may affect the quality of the analysis. The overall similarity among samples were assessed by the euclidean distance between samples (heat-map of sample-to-sample distance) and by principal component analysis. A comparison of gene expression between control and UV-C-treated samples was performed using DESeq2. The Wald test was used to generate *p*-values and log2 fold changes. Genes with an adjusted *p*-value < 0.05 and absolute log2 fold change > 1 were called as differentially expressed genes for each comparison. Volcano plots of the differentially expressed genes were created. Gene ontology (GO) enrichment analysis based on biological process functional categories was performed with ShynyGO v.0.75 with a false discovery rate less than 5% (http://bioinformatics.sdstate.edu/go/).

### Statistics

Statistical analysis was performed using Student’s t test with significance levels of *p* < 0.0001, *p* < 0.001, *p* < 0.01 and *p* < 0.05 for phenotypic characterization and *p*< 0.05 for GA levels and RT-qPCR data.

## Results

### A single UV-C pulse modulates development in Arabidopsis in a biphasic manner

In this study, we investigated the hormonal basis of UV-C induced modulation of Arabidopsis development. In preliminary experiments, we defined the UV-C treatment conditions that induce developmental changes without damaging the plants. To this end, two-week-old Arabidopsis plants were irradiated with 1200 J/m^2^ at a wavelength of 254 nm for 1, 2, 5 or 10 min (Supplementary Fig. S1). Three weeks after treatment, the UV-C-treated plants were phenotypically compared with the untreated control plants (Supplementary Fig. S1, 0 min). A one-minute treatment was found to be most beneficial in terms of plant growth and flower, siliques, and seed production (Supplementary Fig. S1).

Arabidopsis plants treated with UV-C for one minute exhibit a biphasic growth pattern (see Fig. 2a-e). Three days after UV-C treatment, a significant reduction in rosette leaf growth was observed, which lasted for at least ten days after the treatment (Fig. 2c). However, ten days after the treatment, the UV-C-treated plants were taller and had flowered earlier compared to the untreated control plants (Fig. 2a, b, d). The height of the main inflorescence of older plants remained higher in the treated plants compared to untreated plants, increasing by 49% three weeks after the treatment (see Supplementary Fig. S1a).

**Figure 2.**
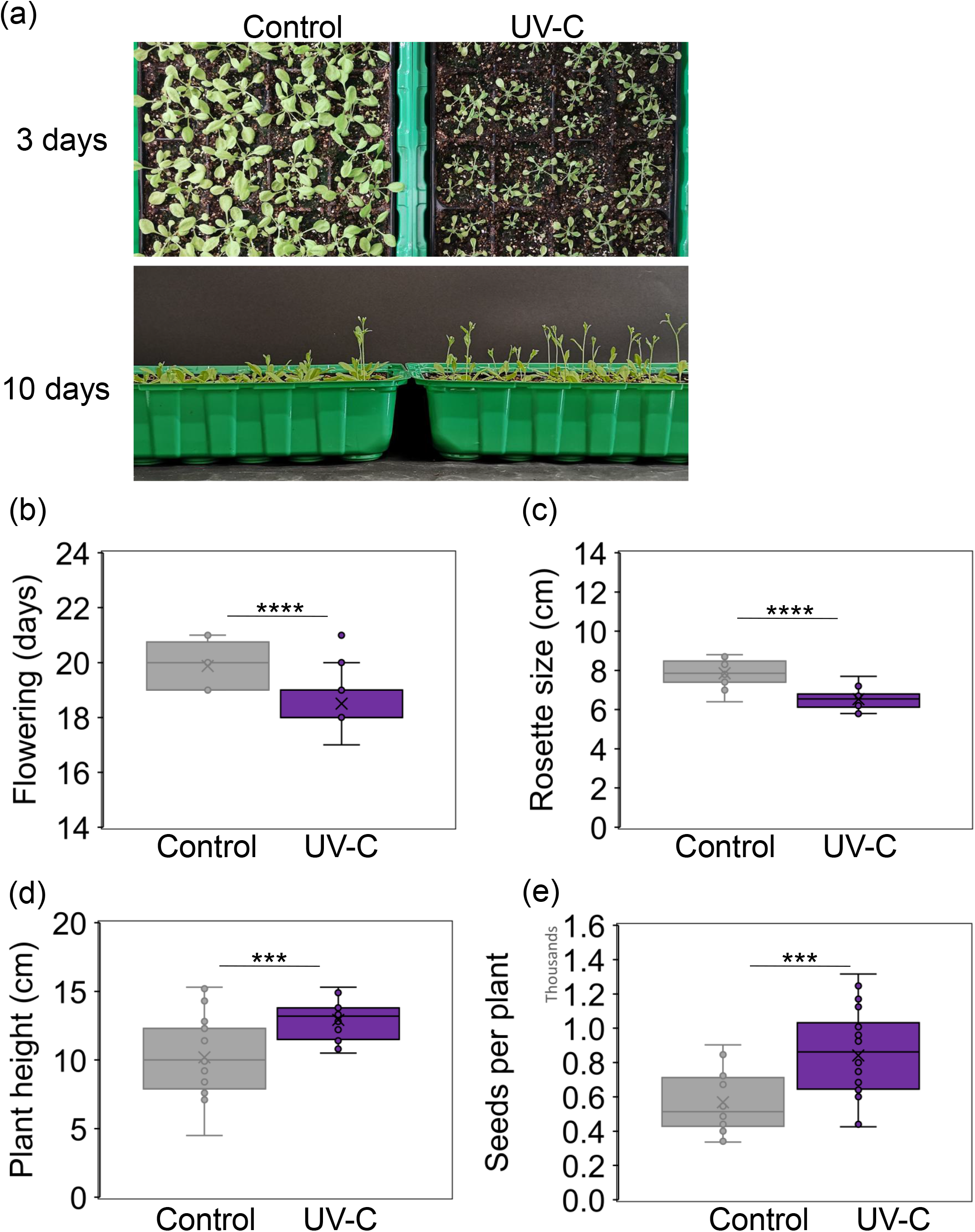
UV-C treatment accelerates development and increases fertility of Arabidopsis plants. The UV-C treatment (0.120 J/cm2 for 1 min at 254 nm) was applied to two-week old plants grown in long days. (a) Phenotypes of control and UV-C treated plants, 3 and 10 days after treatment. (b) Flowering time (days after sowing) of control and UV-C treated plants. (c), (d) Rosette size (rosette leaves diameter) and plant height (height of the main inflorescence) of control and UV-C treated plants, 10 days after treatment. (e) Number of seeds per plant in control and UV-C treated plants, total number of siliques x seed number per silique (estimated from 15 siliques per plant). Data are represented by Box and whisker plots (*n* = 24 plants). Asterisks indicate significant differences between treated and untreated plants (**** *p* < 0.0001, *** *p* < 0.001 Student’s *t*-test).

Previous studies have suggested that stress induces flowering *via* salicylic acid (SA) signalling (Wada and Takeno, 2013; Takeno, 2016). In line with these findings, Martínez *et al*. (2004) reported that accelerated flowering in Arabidopsis following UV-C treatment depended on SA accumulation. The authors described that *nahG* mutant plants, which are unable to accumulate SA, did not show accelerated flowering after UV-C treatment. Furthermore, methyl salicylate was produced in Arabidopsis plants irradiated with UV-C (Yao *et al*., 2011; Xu *et al*., 2016). We therefore applied a one-minute UV-C pulse to the *sid2/N* double mutant, which is deficient in SA biosynthesis (sid2) and accumulation (NahG). Under our growth conditions, *sid2/N* mutant plants respond to the UV-C pulse similarly to wild-type plants. They grow significantly taller and flower significantly earlier after the UV-C treatment, suggesting that SA is not an important regulator of accelerated flowering by UV-C (Supplementary Fig. S2). We have therefore extended our investigations to include gibberellin, a well-known plant hormone that regulates growth and flowering.

### Transcriptome analysis reveals a rapid down-regulation of growth-related hormonal signalling pathways

RNAseq analysis reveals that the number of differentially regulated genes in two-week-old Arabidopsis plants triples at 5, 30, and 120 minutes after one minute of UV-C irradiation, respectively. The number of genes that are upregulated is higher than the number of genes that are downregulated (Supplementary Fig. S3a). Gene Ontology (GO) enrichment analysis using ShinyGO, based on the functional categories of biological processes, revealed an enrichment of signalling pathways associated with defence responses among the genes that were differentially upregulated after UV-C treatment. Conversely, signalling pathways involved in growth were downregulated (see Supplementary Fig. S3b).

Several plant hormone signalling pathways are regulated following the UV-C exposure, including those of abscisic acid, auxin, brassinosteroids, cytokinins, ethylene, jasmonic acid, karrikins, stringolactones, and SA. This suggests that there is rapid hormonal regulation and adaptation of stress responses following UV-C treatment (Supplementary Table S1). Salicylic acid has also been proposed as a regulator of UV-C-induced developmental responses in Arabidopsis, but we were unable to reproduce these results under our growth conditions (see above).

Furthermore, based on published data and our own RNAseq analysis, UV-C treatment also regulates elements of the GA-signalling pathway (Jazayeri et al. 2024, Supplementary Table S1). Several genes encoding enzymes of the GA anabolic pathway were rapidly downregulated after the UV-C pulse, including *ent*-kaurene synthase (KS, (Jazayeri *et al*., 2024)), gibberellin 20-oxidases (*20ox1* and *20ox2*), and gibberellin 3-oxidase1 (*3ox1*, Supplementary Table S1). Conversely, genes of the GA catabolic pathway were upregulated, including gibberellin 2-oxidases (*2ox2, 2ox4, 2ox6*, and *2ox8*). This could indicate that GA signalling plays an important role in UV-C-mediated growth regulation. We have therefore expanded our analysis of GA-related gene expression using qPCR in time-course experiments, as well as analysing endogenous GA levels by GC-MS.

### Rapid downregulation of GA-anabolic, and upregulation of GA-catabolic genes after a one-minute UV-C pulse

Our qPCR experiments confirm that the transcript levels of several genes encoding the initial steps of the GA biosynthetic pathway, including *CPS, KS*, and *KAO2*, are rapidly downregulated after a one-minute UV-C pulse (Fig. 3a). Consistent with the RNAseq results, *20ox1* and *3ox1* transcript levels are lower than in untreated control plants (see Supplementary Fig. S4). Additionally, transcript levels of *2ox6* gene encoding a 2-oxidase of the gibberellin catabolic pathway are rapidly and strongly upregulated until day one, whereas the level of transcripts encoding *2ox1* decrease after UV-C treatment (Fig. 3a). These results could explain the initial retardation of Arabidopsis development observed following UV-C treatment, due to a reduction of early GA-biosynthesis combined with increased GA catabolism *via* 2ox6. We investigated this further by analysing endogenous GA-levels.

**Figure 3.**
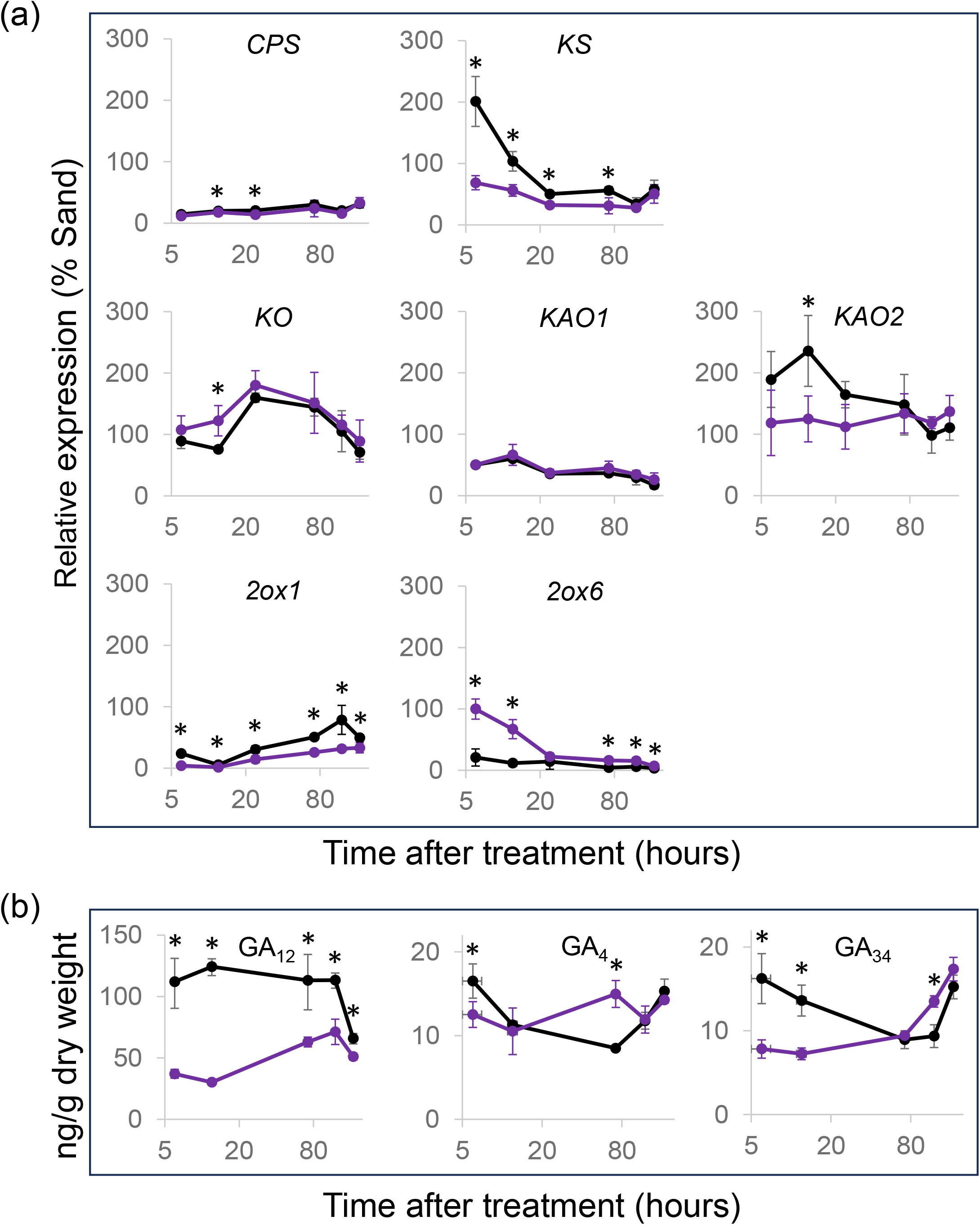
UV-C treatment destabilises Gibberellin homeostasis of Arabidopsis plants. (a) Graphics represent values for relative gene expression measured at different times after UV-C treatment of two-week old Arabidopsis plants grown in long days. (b) Graphics show the endogenous gibberellins measured at different times after UV-C treatment of two-week old Arabidopsis plants grown in long days. (a, b) Black lines, in control plants; violet lines, in UV-C treated plants. Data are shown as means ± CI (*n* = 3). Asterisks indicate significant differences between treated and untreated plants (*p* < 0.05, Student’s *t*-test). Note the log2 scaling of the x-axis.

### Rapid decrease of endogenous GA levels, followed by an increase of GA hormone levels after UV-C treatment

Endogenous levels of several GAs of the biosynthetic pathway decrease rapidly after the UV-C treatment (Fig. 3b, Supplementary Fig. S4b). The levels of the early precursors and catabolic products of the gibberellin pathway, including GA_12_, GA_53_, and GA_110_, remain reduced throughout the entire 7-day period. The levels of the intermediate steps GA_15_ and GA_24_ recover to the levels of the control plants. Initially, the levels of the direct hormone precursor GA_9_, the hormone GA_4_ itself, and the catabolic product GA_34_ decrease, but later increase compared to the levels of untreated control plants (Fig. 3b, Supplemental Fig. 4b). These results suggest that UV-C treatment rapidly inhibits steps of the initial GA-biosynthetic pathway (Fig. 3a), leading to the precursor GA_12_. The subsequent increase in GA_4_ plant hormone levels is consistent with the second phase of elevated growth and development of Arabidopsis as observed after the UV-C treatment. This could indicate higher turnover of the precursors or a decreased overall catabolism (Fig. 3).

### Response of selected GA-mutants to a one-minute UV-C pulse

In order to investigate the contribution of the gibberellin signalling pathway to the UV-C response in Arabidopsis, we tested several mutant lines that exhibit impaired gibberellin signal transduction, including *gdella, kao1, kao2*, and C_19_-*2oxqM*. Under long days (LD), *gdella* plants develop faster than their wild-type Ler counterparts. To optimise the UV-C response, the *gdella* plants were initially grown in short days (SD) for two weeks prior to the UV-C treatment, then left in SD for another week after the treatment. Flower formation was then induced under LD.

Following a one-minute UV-C pulse, the *gdella* mutant exhibits no differences in inflorescence height or flowering time, confirming the involvement of GA signalling in the UV-C response (Fig. 4). Notably, in contrast to the wild-type plants, the number of siliques, their size and the number of seeds are all significantly reduced in the *gdella* mutant treated with a UV-C pulse (Fig. 4b,e-g), suggesting that DELLA plays a protective role in UV-C-induced fertility enhancement.

**Figure 4.**
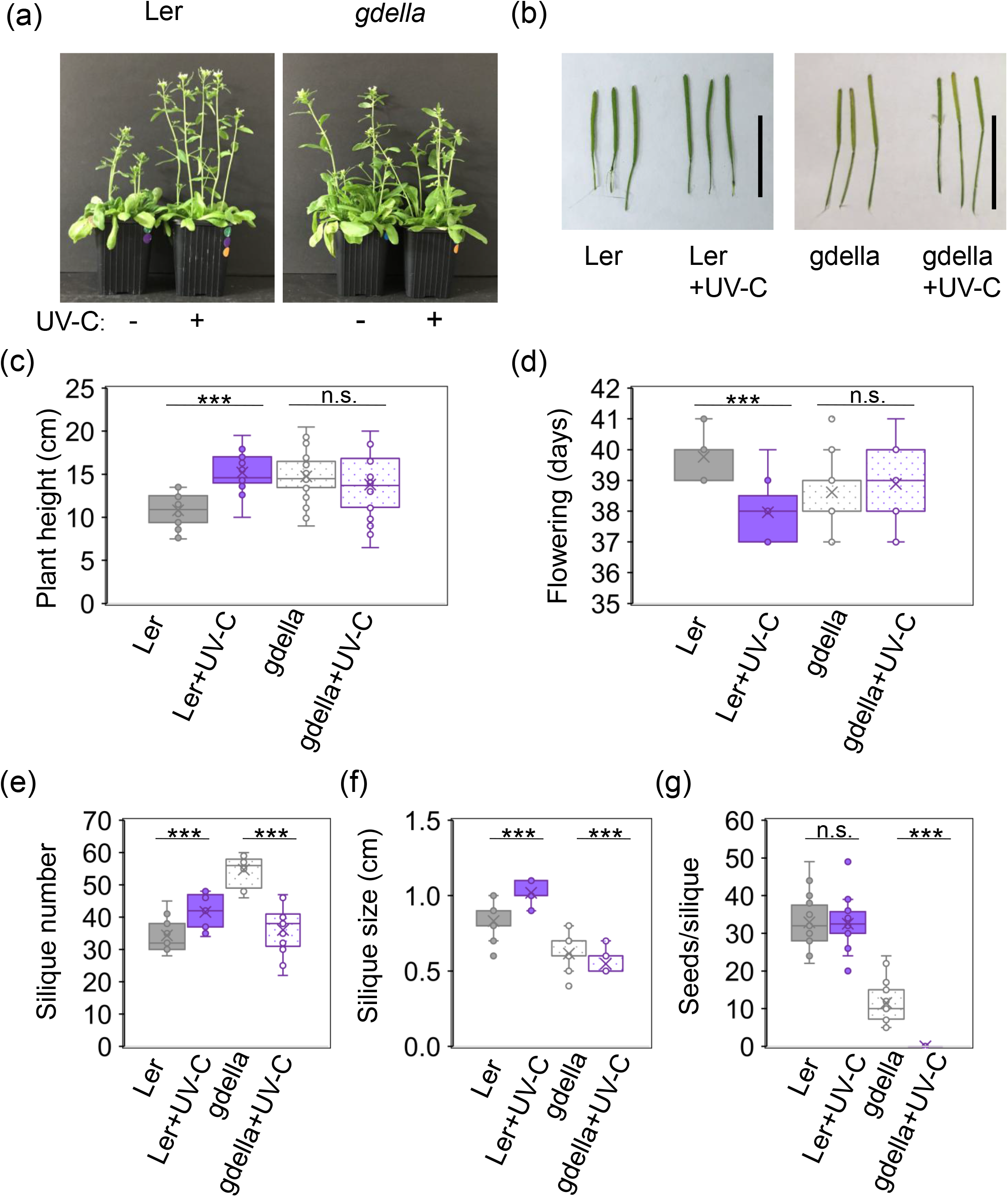
Developmental changes caused by UV-C-treatment in Arabidopsis are regulated by gibberellin. The gibberellin signalling deficient mutant (*gdella)* and respective ecotype (Ler) were grown for 3 weeks in short days before UV-C treatments (1 min). Plants were grown for 1 more week in short days before being moved to long days. (a) Phenotypes of 39-day-old plants untreated or UV-C treated (18 days after treatment). (b) Representative siliques of 60-day-old plants. Bars, represent 1 cm. (c) Plant height, as the height of the main inflorescence of 41-day-old plants (cm). (d) Flowering time (days after sowing). (e) The number of siliques of the main stem (*n* = 15 plants). (f) Silique size (cm, *n* = 50 siliques. (c-g) Data are represented by Box and whisker plots, grey and violet represent untreated and UV-C treated plants, respectively (minimum 26 plants). Asterisks indicate significantly different values, *** *p* < 0.001 and n.s., non-significant difference among samples.

Time-course analyses of transcripts and endogenous gibberellin levels indicate that the early anabolic steps of GA biosynthesis play a key role in the UV-C response (see above). However, mutant plants such as those of *cps, ks*, and *ko* are not suitable for investigating the effects of these biosynthetic steps as they exhibit severe dwarfism alongside an extremely delayed flowering phenotype (Hedden and Sponsel, 2015). Therefore, *kao1* and *kao2* mutants of the early GA biosynthesis pathway were selected that exhibit no obvious phenotype compared to wild-type plants (Regnault *et al*., 2014). Following a one-minute UV-C pulse, both mutants show no significant differences in growth and flowering time. Similar to the *gdella* mutant, the number of siliques decreases compared to untreated mutant plants (Fig. 5), indicating a key role for the initial GA biosynthetic steps in the UV-C response.

**Figure 5.**
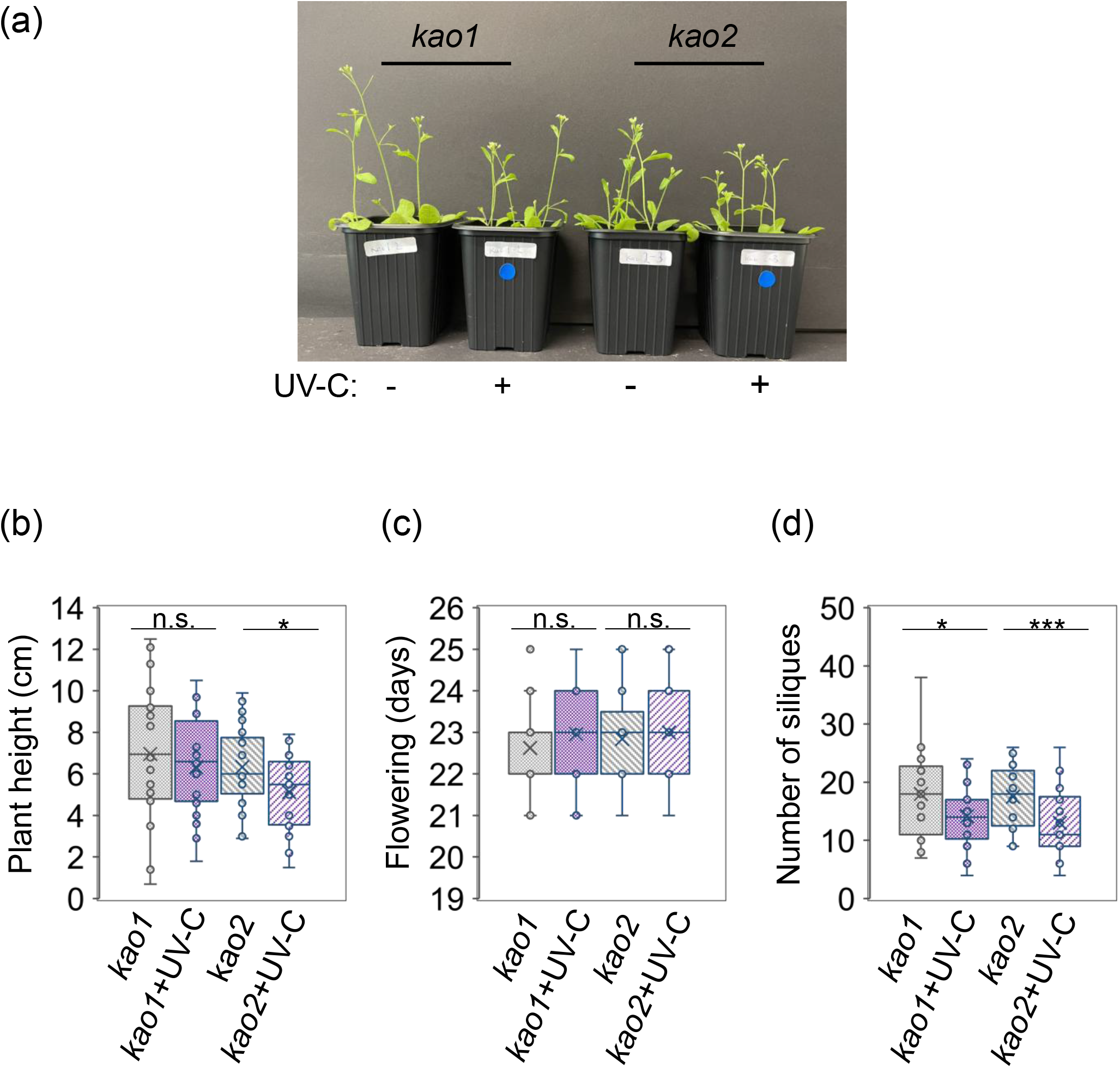
Developmental changes caused by UV-C-treatment in Arabidopsis are regulated by early gibberellin biosynthetic steps. The Arabidopsis gibberellin biosynthetic mutants *kao1* and *kao2* were grown for 2 weeks in long days before UV-C treatments (1 min). (a) Phenotypes of 26-day-old mutant plants, untreated or UV-C treated (12 days after treatment). (b) Plant height, represents the height of the main inflorescence of 26-day-old mutant plants (cm). (c) Flowering time (days after sowing) of untreated and UV-C treated plants. (d) Number of siliques of 35-day-old mutant plants. Data are represented by Box and whisker plots (minimum 25 plants). Asterisks indicate significantly different values, * *p* < 0.05, *** *p* < 0.001, n.s., non-significant difference among samples.

In contrast, the quintuple C_19_-*2oxqM* mutant (*2ox1, 2ox2, 2ox3, 2ox4*, and *2ox6*) responds to the UV-C pulse similarly to wild-type plants: growth increases and flowering time decreases after a one-minute UV-C pulse. This indicates that the five C_19_ 2-oxidases play a limited role in the UV-C regulation of growth and development of Arabidopsis (see Supplementary Fig. S5).

## Discussion

It is known that the application of UV-C light treatments on ornamental plants reduces plant height, increases branching and regulates flowering time. However, the response of plants to UV-C irradiation depends on the dose applied; this can either delay or accelerate flowering (Bridgen, 2016). In Arabidopsis, Martínez *et al*. (2004) reported that UV-C treatment accelerated flowering time *via* SA regulation. Furthermore, UV-C treatment activates plant-to-plant communication, causing UV-C irradiated Arabidopsis plants to produce volatile compounds such as methyl salicylate (MeSA) and methyl jasmonate (MeJA). This results in genome instability not only in the irradiated plants, but also in neighbouring non-irradiated plants (Yao *et al*., 2011; Xu *et al*., 2016).

In our experiments, a one-minute UV-C pulse of 1,200 J/m^2^ was sufficient to alter the growth and development of Arabidopsis in a biphasic manner. Initially, growth was retarded after UV-C treatment, but afterwards, development was enhanced as a form of overcompensation. The plants flowered earlier, were taller, and produced more flowers, siliques, and seeds per plant (Fig. 2, Supplementary Fig. S1). We were unable to confirm regulation *via* SA signalling, as described by Martínez *et al*., (2004), as *sid2/N* mutants responded to the UV-C treatment similarly to wild-type plants. However, the plant hormone GA is known to promote growth and flowering in Arabidopsis by destabilising the growth repressor (DELLA) proteins (Achard *et al*., 2006; Sun, 2008). Arabidopsis plants over-expressing pumpkin GA-biosynthetic genes, *7ox* or *3ox1*, had increased bioactive GA levels and showed accelerated plant development, including earlier flowering time, increased height and a greater number of siliques (Radi *et al*., 2006). Therefore, we hypothesised that altered GA homeostasis might be responsible for the observed changes in Arabidopsis development caused by UV-C treatment.

Immediately following a one-minute UV-C pulse, the transcript levels of several genes encoding different enzymes of gibberellin biosynthetic pathway change. The downregulation of the expression of the initial gibberellin anabolic genes *CPS, KS*, and *KAO2* is consistent with the reduced levels of the gibberellin precursor GA_12_ observed in comparison with untreated control plants. Additionally, the concentrations of GA_12_ by-products GA_53_ and GA_110_ are reduced, suggesting that metabolic pathways branching off from GA_12_, such as 13-hydroxylation or 2ß-hydroxylation, do not contribute to GA_12_ reduction. These results highlight the importance of the initial steps of gibberellin biosynthesis prior to GA_12_ in the UV-C response of Arabidopsis.

Downstream products such as the direct hormone precursor GA_9_, the hormone GA_4_ itself, and its catabolic product GA_34_ show biphasic changes in level compared to controls. Following the UV-C pulse, their levels initially drop, before rising above those of the control plants. This initial drop in GA levels could also have been caused by the reduced availability of the GA_12_ precursor, which is likely to limit the production of downstream GAs. It is also possible that the downstream biosynthetic steps are involved, as we have observed a reduction in transcript levels for the anabolic steps catalysed by 20ox1 and 3ox1. An increase in GA-catabolism (e.g. catalysed by 2ox6), appears unlikely to cause the initial drop in GA_4_ levels, as this would lead to an increase in GA_34_ levels, the opposite of which was observed.

In the second phase, on the third day following UV-C treatment, the GA_4_ levels exceed those of the control plants, before returning to the same level from the fifth day onwards. Catabolism may be involved in this later regulation, as GA_34_ levels increase five days after treatment, when GA_4_ levels return to control plant levels. However, it is unlikely that 2ox1 or 2ox6 are involved; the transcript levels of 2ox6 decrease and of 2ox1 are reduced in this second phase after UV-C treatment, which should lead to a reduction in GA_34_ levels, yet this was not observed. Overall, the two-phase response of the Arabidopsis plants-initially a delay, followed by an acceleration in growth and development - can be linked to an initial decrease, followed by an increase, in GA_4_ hormone levels.

In order to confirm the involvement of the gibberellin signalling pathway and its individual components, we examined various mutants with regard to their response to UV-C irradiation. The *gdella* mutant exhibits no response to UV-C in terms of inflorescence growth or flowering time. This emphasises the significance of the gibberellin signalling pathway in the UV-C response. Furthermore, the *gdella* mutant exhibits reduced fertility following UV-C treatment. This contrasts with the effect observed in wild-type plants, highlighting the protective role of gibberellin signalling during the generative phase.

By contrast, the quintuple C_19_-*2oxqM* mutant responds to a one-minute UV-C pulse in the same way as wild-type plants. Therefore, GA catabolism *via* C_19_ 2-oxidases appears to be insignificant for the UV-C response in Arabidopsis. This also relativises the involvement of the modulation of 2ox1 and 2ox6 at the transcriptional level for the UV-C response, given that their influence on GA hormone levels is limited, as described above.

Similar to the *gdella* mutant, the *kao1* and the *kao2* mutants do not respond to UV-C irradiation with regard to inflorescence growth and flowering time. This demonstrates the importance of the early gibberellin biosynthetic pathway in the UV-C response. Furthermore, as with the *gdella* mutants, the number of siliques in both mutants is reduced after UV-C treatment, indicating its importance in protecting fertility after UV-C treatment. Both, KAO1 and KAO2 have previously been described as functionally redundant in the gibberellin biosynthesis pathway (Regnault et al. 2014). It appears here that both enzymes are necessary for processing the GA-precursors to achieve a full UV-C response.

Our data demonstrate the critical importance of gibberellin signalling, especially of the early stages of gibberellin biosynthesis before GA_12_, in the UV-C response of Arabidopsis. It appears that GA homeostasis is tightly regulated, with outliers towards lower and higher levels that are quickly levelled out. Several key genes were identified as being regulated at the transcriptional level, including *CPS, KS*, and *KAO2*. Feedback and feedforward regulatory mechanisms are likely to modulate the effects of UV-C treatment on the transcription level of several genes, as has been shown, for example for *20ox, 3ox*, and *2ox* genes (Pimenta Lange and Lange, 2006; Hedden, 2020). Additionally, non-transcriptional regulation of the initial steps of gibberellin biosynthesis may contribute to the UV-C responses, akin to the recent findings of (van de Ven *et al*., 2026) regarding regulation of isoprenoid biosynthesis.

The increased defence of plants following UV-C treatment has already been shown numerous times (Urban *et al*., 2018; Ballaré and Austin, 2019; Vàsquez *et al*., 2020; Otake *et al*., 2021; Li *et al*., 2022). Growth-defence trade-offs are widely regarded as a regulatory strategy whereby plants prioritise defence over growth under stressful conditions (He *et al*., 2022). Here, we demonstrate that a brief UV-C pulse can stimulate GA-regulated plant growth and development in Arabidopsis, thereby mitigating the adverse effects of the growth-defence trade-off. This could provide a novel approach for achieving sustainable crop production (Ballaré and Austin, 2019; Li *et al*., 2022).

## Abbreviations

(GA): Gibberellin
(LD): Long days
(SD): Short days
(UV-C): Ultraviolet C

## Acknowledgements

The authors thank Dr. Patrick Achard, Dr. Pepe Cana Quijada and Dr. Andy Philipps for kindly providing Arabidopsis mutant seeds.

## Author contributions

MPL, and TL: conceptualization and methodology; MPL, TL, AC-PM, and JS: investigation and data analysis; MPL, and TL: writing - original draft; MPL, TL, AC-PM, and JS: review & editing; MPL, and TL: supervision; MPL: funding acquisition.

## Data availability

The data that support the findings of this study are available from the corresponding authors upon request.

The RNA-Seq data underlying this article are available in the Gene Expression Omnibus (GEO) under accession GSEXXXX.

## Supplementary data

**Supplementary Table S1**. Gene expression kinetics of selected plant hormone signalling pathways after UV-C treatment.

**Figure S1**. Impact of the duration of UV-C treatment on Arabidopsis development and fertility.

**Figure S2**. Developmental changes caused by UV-C-treatment in Arabidopsis are not regulated by salicylic acid signalling.

**Figure S3**. Transcriptome analysis of early responses to UV-C treatment of Arabidopsis plants.

**Figure S4**. Temporal regulation of the expression of selected GA-oxidase genes after UV-C treatment and endogenous gibberellins measured at different times after UV-C treatment of two-week old Arabidopsis plants.

**Figure S5**. C_19_-GA 2-oxidases do not regulate flowering time and plant height by UV-C-treatment.

